# The CHORD protein CHP-1 regulates EGF receptor trafficking and signaling in *C. elegans* and in human cells

**DOI:** 10.1101/745984

**Authors:** Andrea Haag, Michael Walser, Adrian Henggeler, Alex Hajnal

## Abstract

The intracellular trafficking of growth factor receptors determines the activity of their downstream signaling pathways. The putative co-chaperone CHP-1 acts as a regulator of EGFR trafficking during *C.elegans* vulval development. Loss of *chp-1* causes the retention of the EGFR in the ER and decreased MAPK signaling. CHP-1 functions specifically, as the localization of other receptors is unaltered in *chp-1(lf)* mutants, and inhibiting other co-chaperones does not affect EGFR localization. The role of CHP-1 during EGFR trafficking is conserved in humans. Analogous to *C.elegans*, the response of CHP-1-deficient human cells to EGF stimulation is attenuated, the EGFR accumulates in the ER and ERK2 activity is decreased. Although CHP-1 has been proposed to act as a co-chaperone for HSP90, our data indicate an HSP90-independent function of CHP-1. The identification of CHP-1 as a regulator of EGFR trafficking opens the possibility to identify small molecule chaperone inhibitors targeting the EGFR pathway with increased selectivity.

## Introduction

The generation and maintenance of cellular polarity is essential for the development and homeostasis of organs. Cell polarity governs various processes, such as cell migration, asymmetric cell division and morphogenesis (Bryant & Mostov, 2008). Most of these processes are regulated by extracellular signals, which are received and transduced by specific receptors on the plasma membrane. The intracellular trafficking and subcellular localization of these receptors in polarized epithelial cells profoundly affects their ligand binding capabilities and the activation of the downstream signaling pathways. In particular, the EGFR family of receptor tyrosine kinases, which are activated by a multitude of ligands, play essential roles during the development of most epithelial organs (Citri & Yarden, 2006; Sorkin & Goh, 2009).

In contrast to mammals, *C. elegans* expresses only one EGFR homolog, LET-23, and a single EGF family ligand, LIN-3 (Sundaram, 2006). Thanks to this lack of redundancy, the *C. elegans* EGF/EGFR signaling system is well suited for systematic genetic analysis. LET-23 EGFR signaling controls a variety of developmental processes, including the development of the vulva, the egg-laying organ of the hermaphrodite (Sternberg, 2005). During vulval development, the six vulval precursor cells (VPCs) P3.p to P8.p are induced by an LIN-3 EGF signal from the anchor cell (AC) to differentiate into vulval cells (**Fig. 1A**). The polarized distribution of LET-23 is crucial for the efficient activation of the downstream RAS/MAPK signaling pathway and the induction of the vulval cell fates (Kaech *et al*, 1998; Whitfield *et al*, 1999). P6.p, the VPC closest to the AC, receives the highest dose of LIN-3 and adopts the primary (1°) cell fate. A the same time, P6.p activates via a lateral Delta signal the LIN-12 Notch signaling pathway in its neighbors P5.p and P7.p, which inhibits the 1° and induces the secondary (2°) fate in these VPCs (Sternberg, 2005b; Berset *et al*, 2001). The remaining VPCs P3.p, P4.p & P8.p that receive neither the inductive LIN-3 nor the lateral LIN-12 signal adopt the tertiary (3°) cell fate. The 3° VPCs divide once before fusing to the hypodermis hyp7. Hyperactivation of the EGFR/RAS/MAPK pathway causes more than three VPCs to adopt a vulval cell fate and a multivulva (Muv) phenotype, whereas reduced EGFR/RAS/MAPK signaling results in the induction of fewer than three VPCs and a vulvaless (Vul) phenotype.

**Figure 1.**
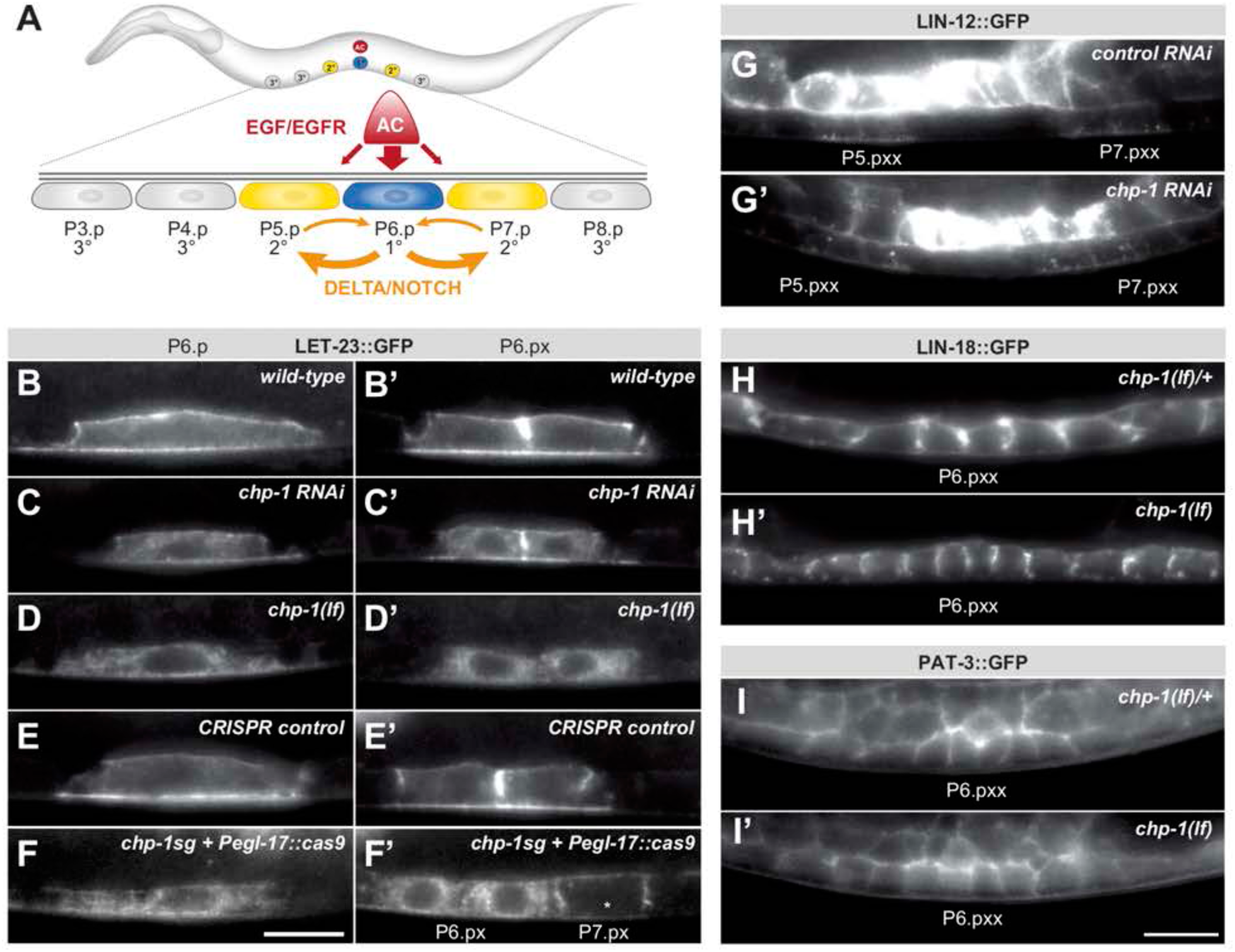
*chp-1* is required for the plasma membrane localization of LET-23::GFP. (**A**) Overview of the EGFR and NOTCH signaling pathways controlling VPC fate determination. (**B**) LET-23::GFP localization in P6.p and (**B’**) the two P6.p daughters (P6.px stage) of a wild-type, (**C, C’**) a *chp-1* RNAi and (**D, D’**) a homozygous *chp-1(tm2277lf)* mutant larva. (**E-F’**) Tissue-specific CRISPR/CAS9 induced deletion of *chp-1* in the 1° vulval cells. (**E,E’**) LET-23::GFP expression in a negative control sibling lacking the transgene and (**F,F’**) an animal carrying the *zhEx558[chp-1sg, egl-17p::cas-9]* transgene. Note in (**F’**) the intracellular mislocalization of LET-23::GFP in the two P6.p descendants, while LET-23::GFP remained localized at the plasma membrane of the P7.p descendants (asterisk). (**G**) LIN-12::GFP localization in a control RNAi and (**G’**) a *chp-1* RNAi treated animals at the Pn.pxx stage. Note the unchanged apical localization of LIN-12::GFP in the 2° P5.p and P7.p descendants (underlined). (**H**) LIN-18::GFP membrane localization in a heterozygous *chp-1(tm2277lf)*/+ and (**H’**) a homozygous *chp-1(tm2277lf)* mutant at the Pn.pxx stage. (**I**) PAT-3::GFP membrane localization in a heterozygous *chp-1(tm2277lf)*/+ and (**I’**) a homozygous *chp-1(tm2277lf)* mutant at the Pn.pxx stage. The 1° P6.p descendants are underlined. At least 20 animals were analyzed for each condition. The scale bars in (**F’**) and (**I’**) are 10 µm.

Thanks to the availability of functional GFP tagged LET-23 reporters and the transparent body, vulval development is an excellent model to observe EGFR trafficking and localization in the epithelial VPCs of living animals (Haag *et al*, 2014). Before vulval induction, LET-23 is equally expressed in all VPCs. During induction, a positive MAP kinase MPK-1 feedback signal up-regulates LET-23 expression in P6.p, which allows this cell to sequester most of the inductive LIN-3 EGF signal (Stetak *et al*, 2006). By contrast, LIN-12 NOTCH signaling in P5.p and P7.p results in the down-regulation of LET-23 and the inhibition of RAS/MAPK signaling (Whitfield *et al*, 1999).

Several factors that regulate RAS/MAPK activity by controlling the sub-cellular localization and trafficking of the LET-23 EGFR in the VPCs have been identified. The basolateral localization of LET-23 by the tripartite LIN-2/CASK, LIN-10/ MINT and LIN-7/VELIS protein complex is necessary for efficient receptor activation, because LIN-3 is secreted by the AC in the somatic gonad facing the basolateral compartment of the VPCs (Kaech *et al*, 1998; Hoskins *et al*, 1996). On the other hand, the ARF GTPase exchange factor AGEF-1 antagonizes via the ARF GTPase and the AP-1 adaptor complex the basolateral localization of LET-23 (Skorobogata *et al*, 2014). A systematic in vivo screen in live *C. elegans* larvae has identified multiple additional regulators of LET-23 localization and signaling (Haag *et al*, 2014). One candidate identified in this screen was *chp-1*, which encodes a Cysteine and Histidine Rich Domain (CHORD) containing protein homologous to human CHORDC1 (also named Morgana) (Brancaccio *et al*, 2003; Ferretti *et al*, 2011). CHORDC1 has been proposed to function as co-chaperone with HSP90, though CHORDC1 may also act independently of HPS90 (Gano & Simon, 2010a).

Here, we show that a loss of *chp-1* function in *C. elegans* leads to the accumulation of LET-23 EGFR in the endoplasmic reticulum (ER) of the VPCs, resulting in a strongly reduced activity of the RAS/MAPK pathway. CHP-1 is specifically required for LET-23 localization, as the secretion of other trans-membrane receptors to the VPC plasma membrane is unchanged in the absence of *chp-1*. Furthermore, we shown that deletion of CHORDC1 in human cells leads to the ER mislocalization of the EGFR and to a loss of EGF-induced filopodia formation. Analogous to *C. elegans chp-1*, deletion of human CHORDC1 does not eliminate but rather attenuates the activation of the MAPK pathway in response to EGF stimulation. We propose that CHP-1 CHORDC1 plays a conserved and specific function during the maturation and membrane secretion of the EGFR.

## Results

### *chp-1* is required for basolateral localization of the EGFR LET-23

The basolateral localization of the LET-23 EGFR in the VPCs of *C. elegans* larvae is necessary for the efficient binding of the LIN-3 EGF ligand secreted by the AC (**Fig. 1A**) (Kaech *et al*, 1998; Whitfield *et al*, 1999). LET-23 is initially secreted to the basolateral membrane compartment, but after ligand-induced receptor endocytosis and transcytosis LET-23 accumulates on the apical cortex of the VPCs (Haag *et al*, 2014). In late L2/early L3 larvae before the VPCs have started dividing, a translational LET-23::GFP reporter was up-regulated in the 1° VPC P6.p, while expression faded in the other, more distal VPCs. Most of the LET-23::GFP protein in P6.p was detected, approximately at equal levels, on the basolateral and apical plasma membranes (**Fig. 1B**). Only a faint and diffuse intracellular LET-23::GFP signal could be observed in the VPCs of wild-type animals. After the first round of VPC divisions, LET-23::GFP continued to be strongly expressed on the plasma membrane of the 1° P6.p descendants (**Fig. 1B’**). A systematic RNA interference screen for genes controlling LET-23 trafficking had previously identified the *chp-1* gene as a regulator of LET-23 localization (Haag *et al*, 2014). *chp-1* encodes a conserved CHORD-containing protein homologous to human CHORDC1/Morgana (Ferretti *et al*, 2011). RNAi against *chp-1* leads to a strong reduction in plasma membrane localization and to the intracellular accumulation of the LET-23::GFP reporter in P6.p and its descendants (**Fig. 1C,C’**). To confirm the RNAi-induced phenotype, we examined LET-23 localization in *chp-1(tm2277)* deletion mutants (*chp-1(lf)*). Since homozygous *chp-1(lf)* mutants are sterile as adults, we analyzed LET-23 EGFR localization in the homozygous offspring of heterozygous *chp-1(lf)*/hT2 balanced mothers. In the following experiments, we compared *chp-1(lf)* homozygous larvae to balanced *chp-1(lf)/hT2* heterozygous control siblings, since LET-23::GFP localization in *chp-1(lf)/+* heterozygotes was indistinguishable from wild-type animals (for example, compare **Fig. 1B** with **Fig. 3A**). Homozygous *chp-1(lf)* L2 and L3 larvae exhibited a completely penetrant intracellular mislocalization of LET-23 EGFR in the VPCs and their descendants (**Fig. 1D,D’**). The distinct plasma membrane signal observed in the VPCs of wild-type animals was absent in *chp-1(lf)* larvae.

In mammalian cells, CHORDC1/Morgana forms a complex with the heat-shock protein 90 (HSP90) and was proposed to function as a co-chaperone for a subset of HSP90 clients (Gano & Simon, 2010a). We therefore tested if a mutation in the *C. elegans hsp-90* homolog *daf-21* affects LET-23 EGFR localization. The *daf-21* reference allele *p673* is sub-viable when the animals are grown at 15°C, but causes larval arrest at higher temperatures (Birnby *et al*, 2000). Even though LET-23::GFP expression in the VPCs of *daf-21(p673)* that had been up-shifted at the L2 stage to 24°C was reduced, most of the LET-23::GFP signal localized at the plasma membrane, and we did not observe an increase in intracellular localization as observed in *chp-1(lf)* mutants (**suppl. Fig. S1A**). Moreover, we examined LET-23::GFP localization after RNAi knock-down of other known HSP90 co-chaperones, such as *cdc-37*, *daf-41* and *sgt-1* (Li *et al*, 2012), but observed no changes in LET-23::GFP localization (**suppl. Fig. S1B-D**).

Taken together, the CHORD-containing protein CHP-1 is required for the plasma membrane localization of the EGFR in the vulval cells independently of the HSP90 chaperone.

### *chp-1* acts cell autonomously in the VPCs

We next tested if *chp-1* acts cell-autonomously in the VPCs by using a tissue-specific CRISPR/Cas9 approach (Shen *et al*, 2014). For this purpose, we expressed the Cas9 endonuclease under control of a 1° VPC-specific *egl-17* promoter fragment (Inoue *et al*, 2002) together with two single-guide (sg) RNAs that target the first exon of *chp-1* and were expressed under control of the ubiquitous *eft-3* promoter (*zhEx558*, see materials and methods). In around 5% of *zhEx558* animals, we observed an intracellular accumulation of LET-23::GFP. In all of these cases, LET-23::GFP was mislocalized only in the 1° VPC P6.p and its descendants (**Fig. 1F,F’**). Control sibling lacking the *zhEx558* array showed a wild-type membrane localization of LET-23::GFP (**Fig. 1E,E’**). The relatively low penetrance of the CRISPR/CAS9-inducd mislocalization phenotype could be due to an inefficient binding of the sgRNAs to the target sequence, to mosaic expression of the extrachromosomal array or to the perdurance of the CHP-1 protein in the VPCs.

These experiments indicated that CHP-1 acts cell-autonomously to regulate LET-23::GFP localization in the VPCs.

### *chp-1* is a specific regulator of LET-23 localization in the VPCs

To investigate if *chp-1* plays a general role in membrane trafficking, we examined the expression pattern of three other trans-membrane receptors expressed in the VPCs; LIN-12 NOTCH (Shaye & Greenwald, 2002), LIN-18 RYK (Inoue *et al*, 2004) and the β-integrin subunit PAT-3 (Hagedorn *et al*, 2009). The translational LIN-12::GFP reporter was expressed on the apical membrane of the VPCs (**Fig. 1G**), while the LIN-18::GFP and PAT-3::GFP translational reporters localized predominantly to the basolateral compartment (**Fig. 1H,I**). *chp-1* RNAi did not alter the apical localization of the LIN-12::GFP reporter (**Fig. 1G’**). Furthermore, the *chp-1(lf)* mutation did not affect the basolateral localization of the LIN-18::GFP or the PAT-3::GFP reporter (**Fig. 1H’,I’**).

Thus, CHP-1 does not play a general role in the apical or basolateral secretion of trans-membrane receptors, but it is rather specifically required for the membrane localization of the EGFR.

### Loss of *chp-1* function causes the accumulation of LET-23 EGFR in the endoplasmic reticulum

The intracellular accumulation of LET-23::GFP in *chp-1(lf)* mutants appeared granular and unevenly structured, while in *chp-1(lf)*/+ control animals, LET-23::GFP was localized predominantly at the plasma membrane of the 1° VPC P6.p and its descendants (**Fig. 2A-C**; note that single mid-sagittal confocal sections through the VPCs are shown in **Fig. 2**, whereas **Figs. 1&3** shows wide-field images of the entire VPCs.) In order to determine the intracellular compartment, in which LET-23::GFP accumulates in *chp-1(lf)* mutants, we generated two reporters that mark the Golgi and the endoplasmic reticulum (ER) of the VPCs, respectively. To label the Golgi apparatus, we expressed an alpha-mannosidase 2A AMAN-2::mCherry fusion protein in the VPCs under control of the pan-epithelial *dlg-1* promoter. AMAN-2 has previously been shown to localize to the Golgi network in the *C. elegans* intestine (Chen *et al*, 2006). In contrast to vertebrate cells that contain one large juxta-nuclear Golgi stack, invertebrate cells contain many small Golgi stacks (Golgi “ministacks”) dispersed throughout the cytoplasm (Ripoche *et al*, 1994). Accordingly, the AMAN-2::mCherry reporter labelled punctate structures scattered throughout the cytoplasm of the VPCs (**Fig. 2A’-F’**). In *chp-1(lf)*/+ animals, LET-23::GFP showed on average 26% co-localization with the AMAN-2::mCherry reporter when analyzed on a voxel per voxel basis in confocal optical sections of the VPCs (**Fig. 2A-C’’** and **suppl. Fig. S2**). In homozygous *chp-1(lf)* mutants, the co-localization with AMAN-2::mCherry was slightly increased to 34.4% (**Fig. 2D-F’’** and **suppl. Fig. S2**). However, the strongest LET-23::GFP signal was detected at the plasma membrane, indicating that only a minor fraction of LET-23::GFP is found in the Golgi apparatus.

**Figure 2.**
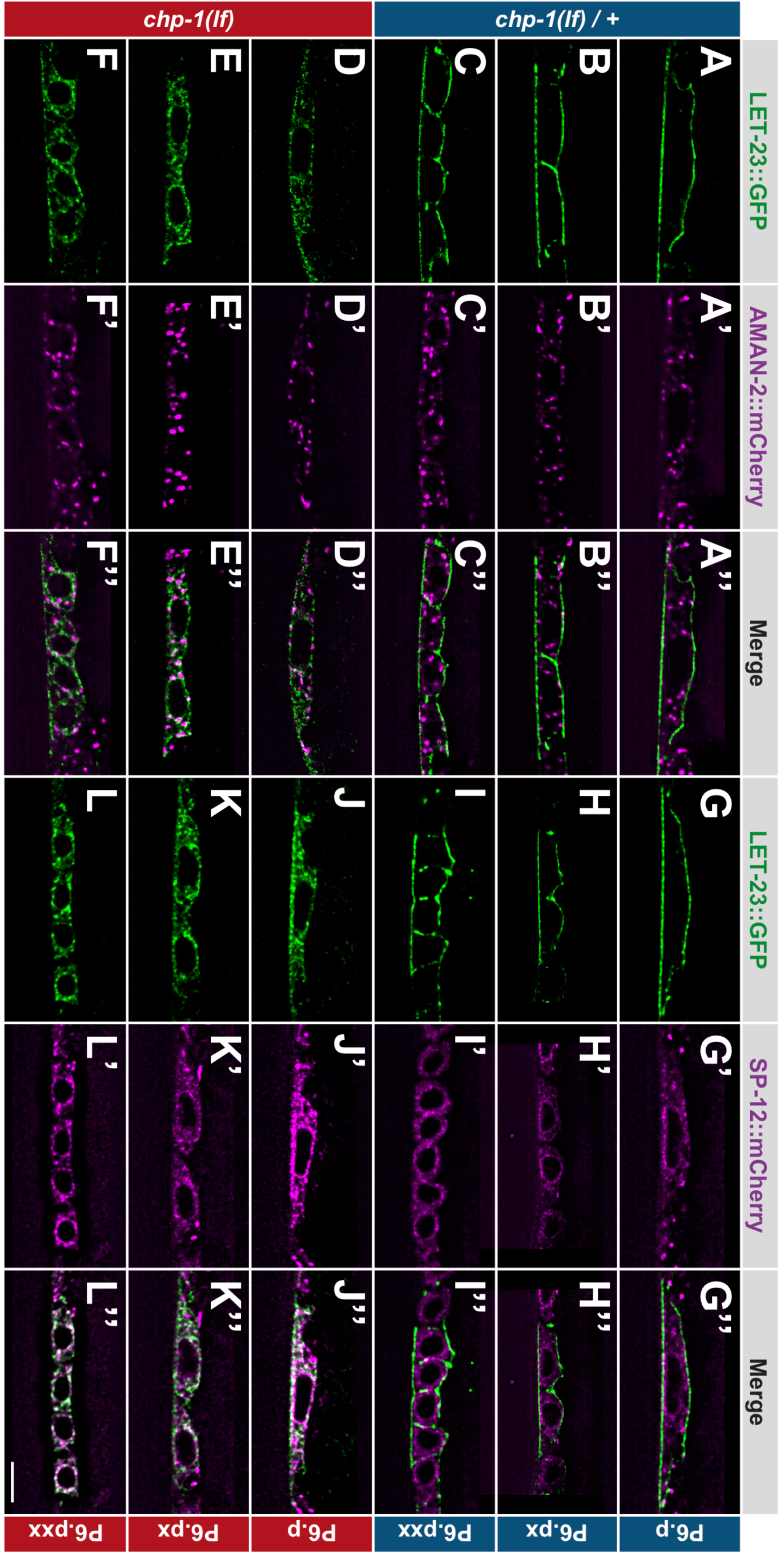
ER mislocalization of LET-23::GFP in *chp-1(lf)* mutants. (**A-C’’**) Localization of LET-23::GFP and the AMAN-2::mCherry Golgi marker in heterozygous *chp-1(tm2277lf)*/+ control siblings and (**D-F’’**) homozygous *chp-1(tm 2277lf)* mutants at the P6.p to P6.pxx stage. (**G-I’’**) Localization of LET-23::GFP and the SP12::mCherry ER marker in heterozygous *chp-1(tm2277lf)*/+ control siblings and (**J-L’’**) homozygous *chp-1(tm 2277lf)* mutants at the P6.p to P6.pxx stage. The individual panels show the different channels of single mid-sagittal confocal sections through P6.p or its descendants. A voxel by voxel quantification of the co-localization between the AMAN-2::mCherry (Golgi) and SP12::mCherry (ER) markers with the LET-23::GFP signal is shown in **suppl. Fig. S2**. The scale bar in (**L’’**) is 10 µm.

To label the ER compartment of the VPCs, we created a reporter consisting of the *dlg-1* promoter driving expression of an *mCherry* tag C-terminally fused to the C34B2.10 gene, which encodes the 12-kDa subunit (SP12) of the *C. elegans* ER signal peptidase complex (Rolls *et al*, 2002). Translational SP12 reporters have previously been shown to localize to a reticular tubular network that extends to the cortex in various cell types of *C. elegans* and resembles the ER architecture in yeast and mammalian cells (Voeltz *et al*, 2002). In confocal sections through the VPCs of *chp-1(lf)*/+ animals, we observed 34% co-localization between the SP12::mCherry and LET-23::GFP reporters, though the strongest LET-23::GFP signal intensity was found at the cell cortex where no SP12::mCherry was detected (**Fig. 2G-I’’** and **suppl. Fig. S2**). By contrast, in homozygous *chp-1(lf)* mutants we observed a strong overlap between the LET-23::GFP and SP12::mCherry signals inside the cells, resulting in 64% co-localization between the two reporters (**Fig. 2J-L’’** and **suppl. Fig. S2**).

Therefore, the loss of *chp-1* function leads to the intracellular retention of LET-23 mainly in the ER.

### ER mislocalization of LET-23 in *chp-1(lf)* mutants does not activate the unfolded protein response pathway

Since LET-23::GFP accumulated mainly in the ER compartment of *chp-1(lf)* mutants, we tested if loss of *chp-1* function causes ER stress triggering the unfolded protein response (UPR) pathway. The *hsp-4* gene encodes a homolog of the mammalian Grp78/BiP protein that is upregulated upon ER stress via the XBP-1 transcription factor and the IRE-1 kinase/endoribonuclease (Calfon *et al*, 2002). The expression of an *hsp-4::gfp* reporter thus serves as an in vivo readout for the UPR (Taylor & Dillin, 2013). As a positive control, we treated animals with tunicamycin, which induces UPR by blocking the formation of N-acetylglucosamine lipid intermediates necessary for the glycosylation of newly synthesized proteins in the ER (Taylor & Dillin, 2013). Untreated *chp-1(lf)* mutants did not show elevated *hsp-4*::GFP expression when compared to the wild-type (**suppl. Fig. S3A,C,E**). A 4 hour exposure of young adult wild-type animals to 25µg/ml tunicamycin caused an approximately 8-fold increase in *hsp-4*::GFP fluorescence intensity (**suppl. Fig. S3B,E**). Interestingly, tunicamycin-treated *chp-1(lf)* mutants exhibited a stronger induction of *hsp-4*::GFP expression (**suppl. Fig. S3D,E**).

Taken together, the intracellular accumulation of LET-23 in the *chp-1(lf)* mutants does not activate the UPR pathway under standard conditions. However, *chp-1(lf)* mutants are slightly hypersensitive to ER stress induced by tunicamycin-treatment.

### Intracellular LET-23 EGFR accumulation in *chp-1(lf)* mutants is ligand-independent

Binding of the LIN-3 EGF ligand to the LET-23 EGFR on the basolateral cortex of the VPCs induces rapid receptor endocytosis (Haag *et al*, 2014). The endocytosed LET-23 can be recycled to the basolateral compartment, transported via transcytosis to the apical membrane compartment or undergo lysosomal degradation (Whitfield *et al*, 1999; Stetak *et al*, 2006). In *lin-3(e1417)* mutants, in which LIN-3 expression is specifically reduced in the AC (Hwang & Sternberg, 2004), LET-23::GFP accumulated on the basolateral cortex of the VPCs, while the apical LET-23::GFP signal was reduced (**Fig. 3C**) (Haag *et al*, 2014). We therefore examined whether the intracellular accumulation of LET-23 in *chp-1(lf)* mutants depends on ligand-induced receptor endocytosis. In *lin-3(e1417)*; *chp-1(lf)* double mutants, we observed the same intracellular accumulation of LET-23::GFP as in *chp-1(lf)* single mutants (**Fig. 3B,D**). We further tested if a global reduction in endocytosis alters LET-23::GFP localization in *chp-1(lf)* mutants. For this purpose, we performed RNAi against *rab-5*, which encodes a small GTPase that functions as a key regulator of early endosome formation and was previously shown to control LET-23 trafficking (Skorobogata & Rocheleau, 2012). Similar to the *lin-3(e1417)* mutation, *rab-5i* did not prevent the intracellular accumulation of LET-23::GFP in *chp-1(lf)* mutants, indicating that *chp-1(lf)* does not cause the accumulation of LET-23 in the endocytic compartment (**Fig. 3E,F**).

**Figure 3.**
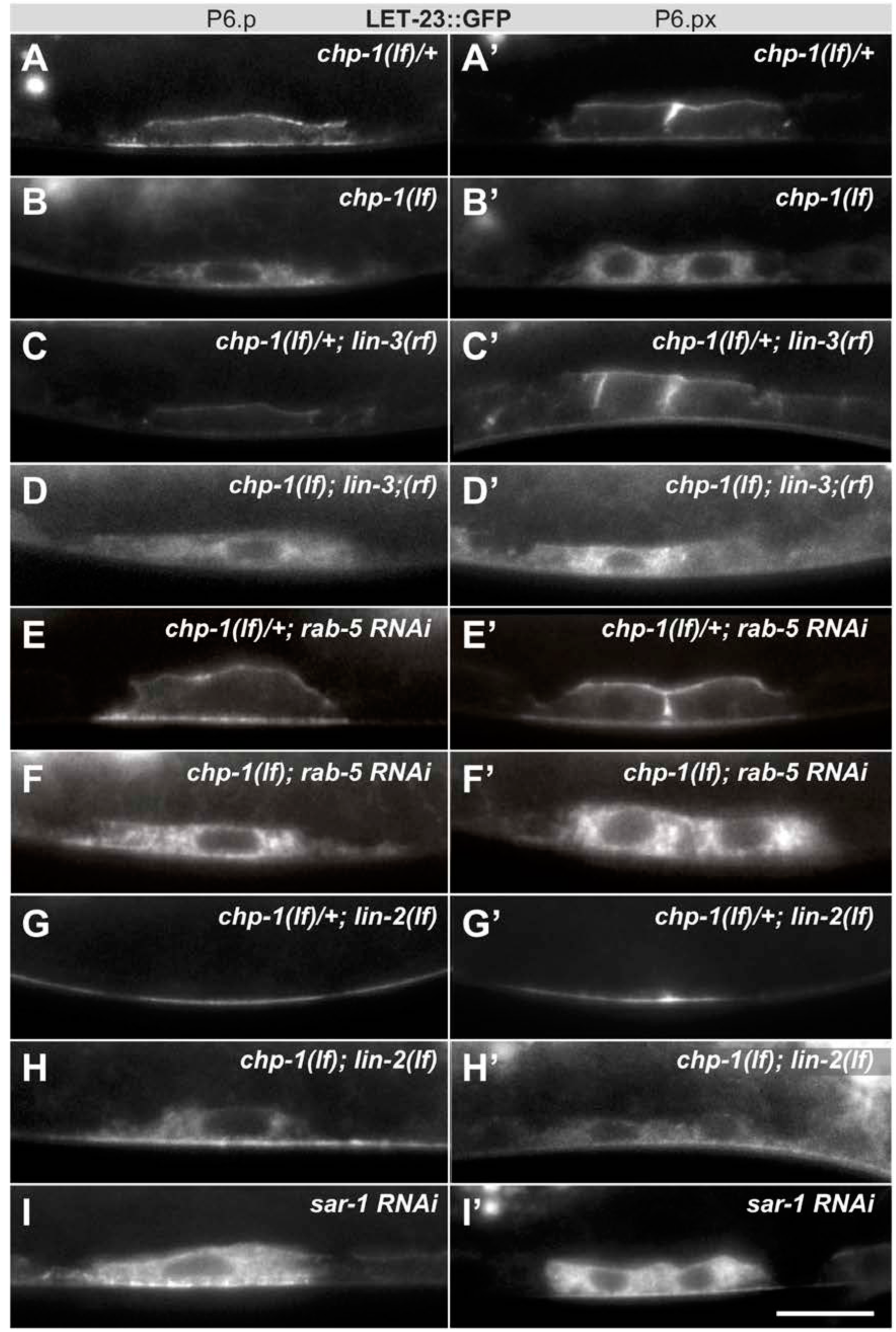
Intracellular mislocalization of LET-23::GFP in *chp-1(lf)* mutants is ligand independent. (A, A’) Localization of LET-23::GFP in heterozygous *chp-1(tm2277lf)*/+ control siblings, (B, B’) homozygous *chp-1(tm 2277lf)* mutants, (C, C’) *lin-3(e1417rf)* single mutants, and (D, D’) in *chp-1(tm2277lf); lin-3(e1417rf)* double mutants at the P6.p and P6.px stage. (E,E’) LET-23::GFP localization in *rab-5* RNAi treated heterozygous *chp-1(tm2277lf)*/+ controls and (F, F’) homozygous *chp-1(tm 2277lf)* mutants. (G, G’) Apical mislocalization of LET-23::GFP in *lin-2(n397lf)* single mutants and (H, H’) intracellular LET-23::GFP localization in *chp-1(tm2277lf); lin-2(n397lf)* double mutants. (I,I’) *sar-1* RNAi causes the same intracellular accumulation of LET-23::GFP as *chp-1(lf)*. At least 20 animals were analyzed for each condition. The scale bar in (I’) is 10 µm.

Next, we asked whether *chp-1* acts at the level of the tripartite LIN-2/LIN-7/LIN-10 complex, which is required for the basolateral retention of LET-23 and facilitates ligand binding (Kaech *et al*, 1998; Whitfield *et al*, 1999). Mutations in *lin-2*, *lin-7* or *lin-10* cause a penetrant vulvaless (Vul) phenotype because LET-23 is mislocalized to the apical membrane compartment, where it cannot bind to LIN-3 secreted by the AC on the basal side of the VPCs. In *lin-2(lf)* single mutants, the LET-23::GFP signal was detected almost exclusively on the apical membranes of the VPCs (**Fig. 3G**). In *chp-1(lf)*; *lin-2(lf)* double mutants a major fraction of the LET-23::GFP signal was found in the intracellular compartment, similar to *chp-1(lf)* single mutants (**Fig. 3H**). However, we did observe a faint LET-23::GFP signal on the apical cortex in *chp-1(lf)*; *lin-2(lf)* double mutants, indicating that a fraction of LET-23::GFP can be secreted to the apical plasma membrane in the absence of *chp-1*. Finally, RNAi against *sar-1*, which encodes a small GTP binding protein required for ER to Golgi transport, caused the same intracellular mislocalization of LET-23::GFP as observed in *chp-1(lf)* mutants (**Fig. 3I**).

Taken together, we conclude that CHP-1 does not regulate the ligand-induced endocytosis or basolateral retention of LET-23, but rather the secretion of the receptor from the ER to the plasma membrane.

### *chp-1* is a positive regulator of EGFR/RAS/MAPK signaling in the VPCs

The basolateral membrane localization of LET-23 is necessary for efficient ligand binding and activation of the downstream RAS/MAPK signaling pathway in the VPCs (Kaech *et al*, 1998; Hoskins *et al*, 1996; Whitfield *et al*, 1999). To quantify the output of the RAS/MAPK pathway in the VPCs, we analyzed the expression of a transcriptional P*_egl-17_::cfp* reporter as a marker for the 1° cell fate (Burdine *et al*, 1998). *egl-17* encodes a fibroblast growth factor (FGF)-like protein, which is up-regulated by RAS/MAPK signaling in the 1° VPC (P6.p) and its descendants until the late L3 stage. P*_egl-17_::cfp* expression in the VPCs of *chp-1(lf)* mutants at the Pn.pxx stage was decreased around five-fold when compared to wild-type larvae at the same stage (**Fig. 4A,B,E**). We also examined the activity of the lateral NOTCH signaling pathway using a transcriptional P*_lip-1_::gfp* reporter that is up-regulated in 2° VPC in response to LIN-12 NOTCH activation (Berset *et al*, 2001). Expression of the P*_lip-1_*::*gfp* reporter was unchanged in *chp-1(lf)* mutants (**Fig. 4C,D**). Thus, CHP-1 acts as a positive regulator of RAS/MAPK signaling, while the activity of the lateral NOTCH pathway is not affected by *chp-1(lf)*.

**Figure 4.**
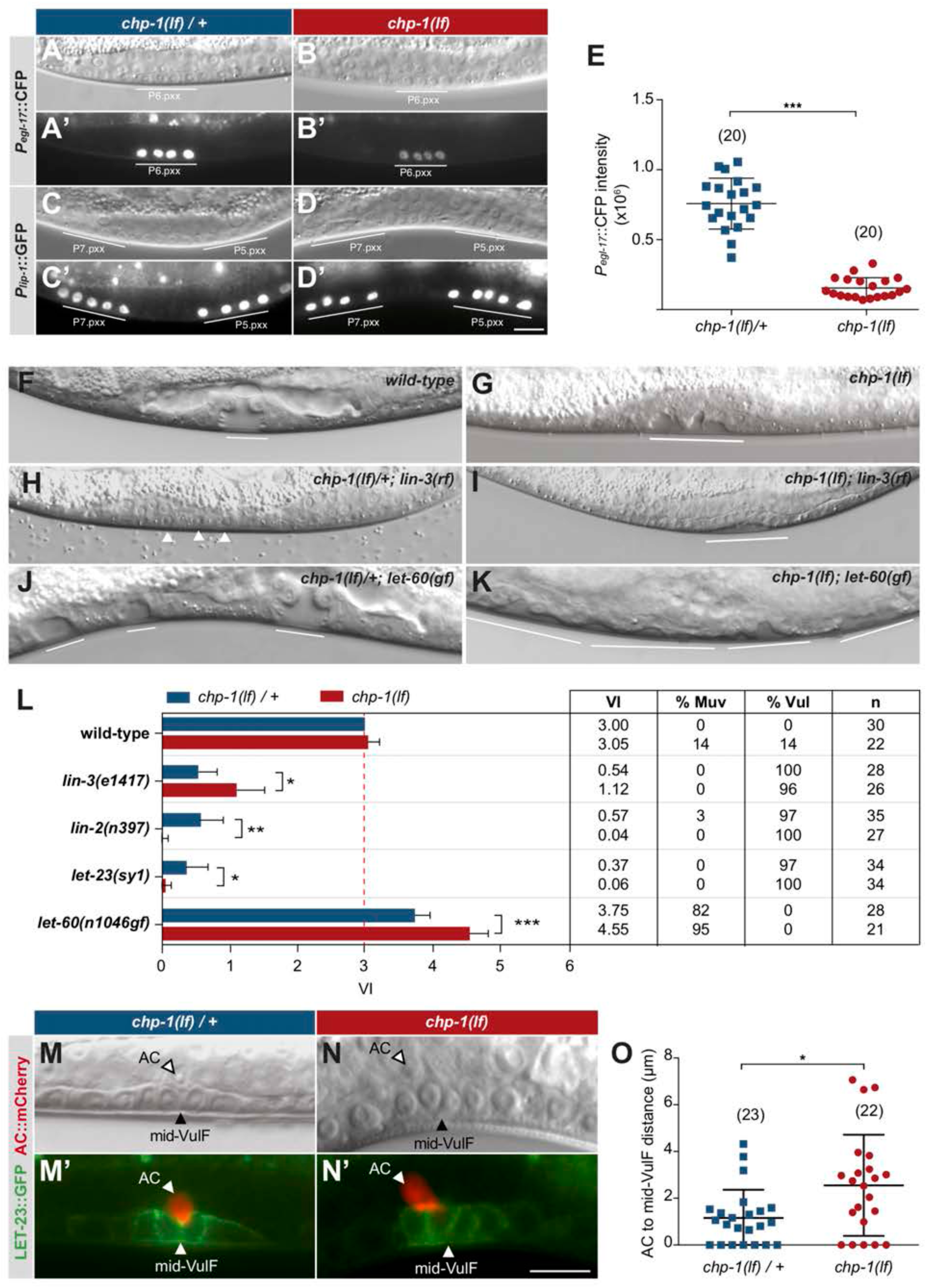
CHP-1 positively regulates EGFR/RAS/MAPK signaling in the VPCs. (**A, A’**) Expression of the 1° cell fate reporter P*egl-17*::CFP in heterozygous *chp-1(tm2277lf)*/+ and (**B, B’**) homozygous *chp-1(lf)* mutant at the Pn.pxx stage. The top panels show Nomarski images of the differentiating VPCs and the bottom panel the reporter expression taken with identical exposure settings. (**C, C’**) Expression of the 2° cell fate reporter P*lip-1*::GFP in heterozygous *chp-1(tm2277lf)*/+ and (**D, D’**) homozygous *chp-1(lf)* mutant at the Pn.pxx stage. The scale bar in (**D’**) is 10 µm. (**E**) Quantification of the P*egl-17*::CFP fluorescence intensity in the 1° VPCs at the Pn.pxx stage. The p-value was calculated by a two-tailed t-test for independent samples. The numbers of animals quantified are indicated in brackets. (**F**) Nomarski images of the vulval morphology in wild-type, (**G**) homozygous *chp-1(tm2277lf)*, (**H**) heterozygous *chp-1(tm2277lf)*/+; *lin-3(e1417rf)*, (**I**) homozygous *chp-1(tm2277lf)*; *lin-3(e1417rf)*, (**J**) heterozygous *chp-1(tm2277lf)*/+; *let-60(n1046gf)* and **(K**) homozygous *chp-1(tm2277lf)*; *let-60(n1046gf)* L4 larvae. The descendants of induced VPCs forming an invagination are underlined and the arrowheads in (**H**) point at the nuclei of uninduced VPCs. (**L**) Quantification of the vulval induction index (VI) for the indicated genotypes. The table to the right shows the absolute mean VI, the percentage of animals with a Muv (VI>3) and a Vul (VI<3) phenotype and the number of animals scored (n) for each genotype. Error bars indicating the 95% confidence intervals and p-values were calculated by Bootstrapping with a resampling size of 1000, as described in Maxeiner *et al*. (2019) (p < 0.05 = * p < 0.01 = ** and p < 0.001 = ***). (**M, M’**) AC to 1° VPC alignment at the Pn.pxx stage in heterozygous *chp-1(tm2277lf)/+* and (**N, N’**) homozygous *chp-1(tm2277lf)* mutants. The top panels show Nomarski images and the bottom panels the expression of the LET-23::GFP reporter in green and the *qyIs23[Pcdh-3:: PLC∂PH::mCherry]* reporter labelling the AC in red. The scale bar in (**N’**) is 10 µm. (**O**) Quantification of the AC to VulF midline distance. The p-value was calculated by a two-tailed t-test for independent samples, and the numbers of animals scored are indicated in brackets.

Despite the strong reduction in RAS/MAPK reporter expression, the VPCs in *chp-1(lf)* mutants were induced to proliferate and differentiate into vulval cells. In most *chp-1(lf)* single mutants, the three proximal VPCs P5.p through P7.p differentiated, as in wild-type animals (**Fig. 4F,G**). However, the vulval invagination formed by the descendants of the induced VPCs had an abnormal shape and the vulval cells formed two separate invaginations, indicating that CHP-1 performs additional functions during vulval morphogenesis (**Fig. 4G**). To quantify vulval induction, we determined the vulval induction index (VI) by counting the average number of VPCs per animals that were induced to differentiate (Schmid *et al*, 2015). A VI of 3 indicates wild-type differentiation, while a VI>3 signifies over- and a VI<3 under-induction. The VI thus serves as a quantitative readout to examine genetic interactions between signaling pathway components. Most *chp-1(lf)* mutants showed a VI of 3, though we observed over- as well as under-induced animals (**Fig. 4L**).

To investigate the interaction between *chp-1* and the EGFR/RAS/MAPK signaling pathway, we constructed double mutants between *chp-1(lf)* and core EGFR/RAS/MAPK pathway components. The *lin-3(e1417rf)* allele caused a strong reduction in the VI of *chp-1(lf)* mutants, approximately to the level of *lin-3(e1417)* single mutants (**Fig. 4H,I,L**) (Hwang & Sternberg, 2004). This indicates that the VPCs in *chp-1(lf)* mutants are at least partially sensitive to the inductive AC signal. Moreover, double mutants between *chp-1(lf)* and *let-23 egfr(sy1)* or *lin-2(lf)* exhibited a significantly stronger Vul phenotype than *let-23(sy1)* or *lin-2(lf)* single mutants (**Fig. 4L**). The *let-23*(*sy1)* allele specifically prevents the interaction of LET-23 EGFR with the LIN-2/LIN-7/LIN-10 receptor localization complex and causes a similar apical receptor mislocalization and partially penetrant Vul phenotype as the *lin-2(lf)* mutation (Kaech *et al*, 1998; Whitfield *et al*, 1999). Thus, *chp-1(lf)* enhanced the Vul phenotype caused by apical receptor mislocalization. Interestingly, the Muv phenotype caused by the *n1046* gain-of-function (*gf*) mutation in the *let-60 ras* gene (Beitel *et al*, 1990) was significantly enhanced by *chp-1(lf)* (**Fig. 4J-L**). This apparent paradox may be explained by reduced LIN-3 EGF sequestering in the proximal VPC P6.p of *chp-1*(*lf*) mutants. Similar to the apical mislocalization in *lin-2(lf)* mutants, the intracellular mislocalization of LET-23 in *chp-1(lf)* mutants likely results in decreased ligand binding by P6.p, allowing in the diffusion of more LIN-3 signal to distal VPCs and their induction in the hyper-sensitive *let-60(n1046gf)* background (Hajnal *et al*, 1997).

In summary, our genetic analysis indicated that *chp-1* positively regulates EGFR/RAS/MAPK signaling in the VPCs. However, the VPCs in *chp-1(lf)* mutants can differentiate into vulval cells because they remain partially sensitive to the inductive LIN-3 EGF signal.

### CHP-1 is necessary for the precise AC to P6.p alignment

Besides VPC fate specification, LIN-3 to LET-23 signaling is also required for the proper alignment between the AC and the 1° VPC P6.p (Grimbert *et al*, 2016). In wild-type L2 stage larvae, the relative position between the AC and VPCs is highly variable. However, by the early L3 stage the 1° VPC P6.p has migrated towards the AC such that the AC and P6.p are precisely aligned with each other. Thereafter, the VPCs begin to proliferate and the AC remains aligned with the 1° VPC descendants. In wild-type mid-L3 larvae after the VPCs had undergone two rounds of cell divisions, the AC was located at the vulval midline above the two inner 1° P6.p descendants (the VulF cells) (**Fig. 4M,M’**). By contrast, in *chp-1(lf)* mutants the AC was often misplaced and occasionally located between VulF and VulE (**Fig. 4N,N**). To quantify the AC to VulF alignment, we measured the distance between the AC nucleus and the midpoint between the two VulF cells at the Pn.pxx stage. In most *chp-1(lf)/+* heterozygous control animals, the AC to mid-VulF distance was around 1 µm or smaller, while most *chp-1(lf)* mutants exhibited a distance greater than 1 µm (**Fig. 4O**).

Thus, in addition to VPC fate specification *chp-1* is also required for the precise alignment between the AC and the VPCs mediated by LIN-3/LET-23 signaling.

### Human CHORDC1 is required for filopodia formation and sustained ERK1/2 activation by EGF

To examine if the role of CHP-1 in controlling EGFR localization and signaling is conserved in mammalian cells, we performed a CRISPR/Cas9-mediated knock-out of the mammalian *chp-1* homolog CHORDC1 in cultured cells. We used the human vulva epidermoid carcinoma cell line A431 because the cells express high levels of wild-type EGFR and respond strongly to EGF stimulation (Van de Vijver *et al*, 1991). A431 cells were transduced with lentiviral particles that deliver Cas9 and two sgRNAs targeting the first exon of CHORDC1 (see materials and methods). As negative control, cells were transduced with a lentivirus delivering a scrambled sgRNA. After bulk puromycin selection to eliminate uninfected cells, two cell populations were generated, subsequently termed A431 KO and A431 control cells, respectively. Western blot analysis revealed a 96% reduction of CHORDC1 protein levels in A431 KO cells (**Fig. 5A**).

**Figure 5.**
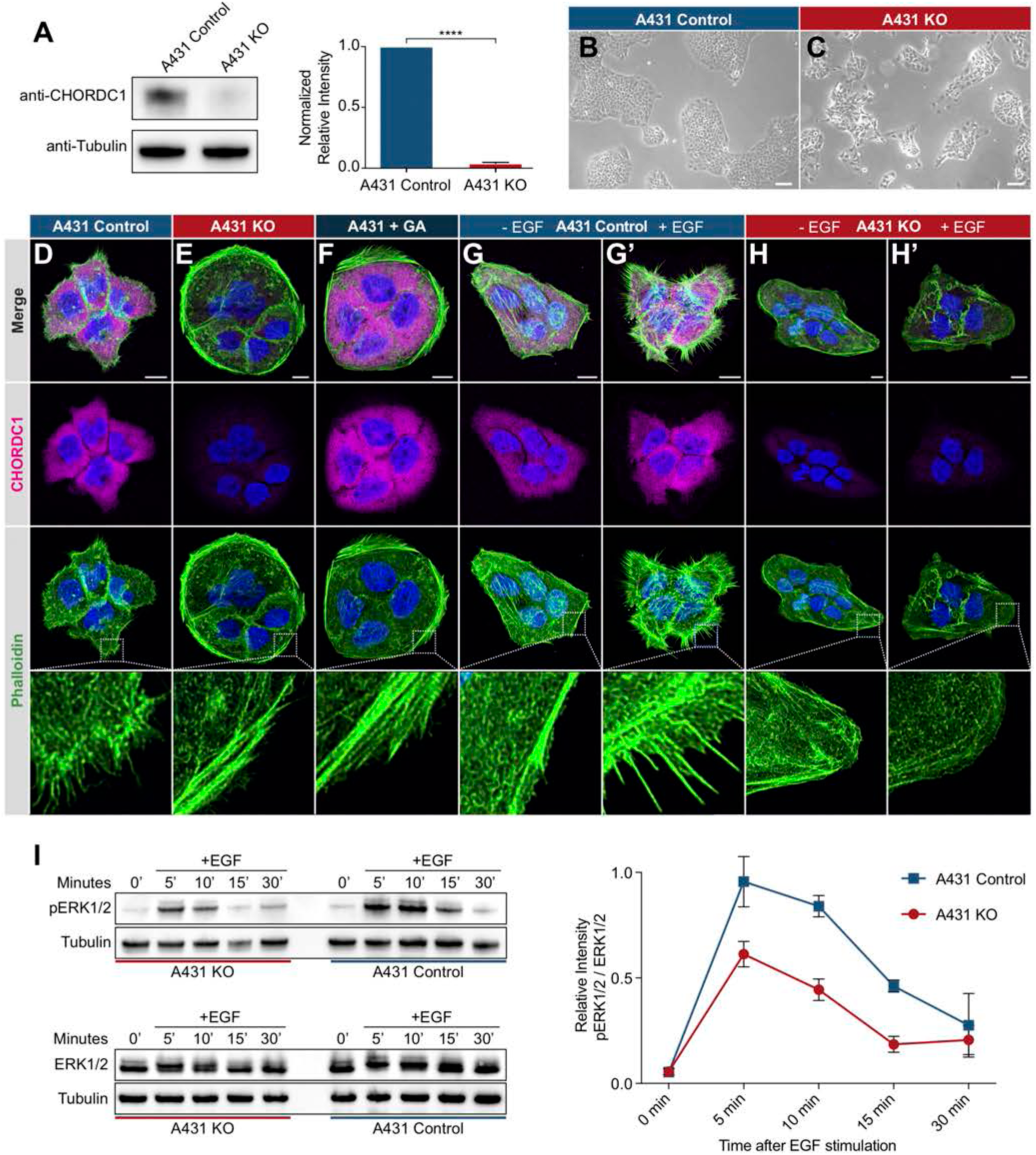
CHORDC1 is required for EGF-induced filopodia formation and sustained ERK activation in A431 cells. (**A**) Quantification of CHORDC1 protein levels in A431 control and KO cells by Western blot analysis (average of eleven biological replicates). The bar graph shows the normalized averaged relative intensities ± SEM. The p-value was calculated using a two-tailed t-test for independent samples. (**B**) Phase contrast images of A431 control and (**C**) A431 KO cells 12 days post lentiviral transduction. The scale bars are 100 µm. (**D**) Immunofluorescence staining of A431 control cells with antibodies recognizing CHORDC1 (magenta), fluorescently-labeled phalloidin (green) and DAPI (blue), (**E**) A431 KO cells, and (**F**) A431 cells treated for 24 hours with 1 µM geldanamycin (GA). (**G**, **H**) Control- and KO cells fixed after 16 hours of serum starvation, and (**G’**, **H’**) 10 minutes after stimulation with 100 ng/ml human EGF. The scale bars are 10 µm. The bottom row shows higher magnifications of the cortical regions outlined by the dashed squares. (**I**) Total protein lysates of serum starved cells that had been stimulated with 100 ng/ml human EGF for the indicated times (in minutes) were analyzed on Western blots with antibodies against phospho-ERK1/2 and total ERK1/2. The graph to the right shows the relative phospho-ERK1/2 signals normalized to the total ERK1/2 levels at each time point. The data shown represent the average ratios obtained in three biological replicates. Error bars indicate the SEM.

A431 KO cells displayed a reduced growth rate and a tendency to grow in smaller, scattered patches when compared to A431 control cells (**Fig. 5B,C**). Furthermore, loss of CHORDC1 resulted in cell lethality approximately 16 days post lentiviral transduction, which made it impossible to establish A431 KO lines from single cell clones. Therefore, the following experiments were performed with populations of A431 KO and control cells 10 to 14 days post lentiviral transduction. To further characterize the morphological defects of A431 KO cells, we visualized the actin cytoskeleton by phalloidin staining. The numerous F-actin rich filopodia protruding from the plasma membrane of A431 control cells were absent in A431 KO cells (**Fig. 5D,E**). Instead, A431 KO cells contained densely packed cortical F-actin filaments arranged in a circumferential manner. A similar phenotypic switch was observed after treatment of A431 cells with the Hsp90 inhibitor geldanamycin (GA) (**Fig. 5F**) (Ahsan *et al*, 2012; Gano & Simon, 2010b).

Since EGFR signaling induces the remodeling of the actin cytoskeleton in migratory cells (Appert-Collin *et al*, 2015), we tested if the absence of filopodia in A431 KO cells might be due to reduced EGFR signaling. Serum starvation of A431 control cells caused a strong reduction in filopodia formation (**Fig. 5G**), but stimulation with EGF induced the reappearance of actin-rich filopodia within 10 minutes (**Fig. 5G’**). By contrast, EGF stimulation of serum starved A431 KO cells did not induce filopodia formation (**Fig. 5H, H’**).

To directly measure the activity of the EGFR/RAS/MAPK pathway, we quantified ERK1/2 activity after EGF stimulation of serum starved cells using a phospho-ERK1/2 specific antibody to probe Western blots of total cell lysates (Gabay *et al*, 1997). In A431 control cells, phospho-ERK1/2 levels reached the maximum levels 5 minutes after EGF stimulation and declined almost to baseline levels within 30 minutes (**Fig. 5I**). The total ERK1/2 levels did not change during the EGF stimulation. By contrast, phospho-ERK1/2 levels in A431 KO cells increased to around half of the maximal levels observed in A431 KO cells and decreased more rapidly.

Taken together, our results show that the human CHP-1 homolog CHORDC1 is required for EGF-induced filopodia formation and sustained ERK1/2 activation in A431 cells. Analogous to the results obtained for *C. elegans chp-1*, loss of CHORDC1 function does not eliminate but rather attenuates EGFR signaling in A431 cells.

### CHORDC1 controls the sub-cellular localization and stability of the EGFR

Since *chp-1* is required for the membrane localization of the LET-23 EGFR in *C. elegans*, we investigated if CHORDC1 also regulates EGFR localization in A431 cells. We analyzed receptor localization by immunofluorescence staining of fixed cells with an antibody against the extracellular domain of the EGFR. In A431 control cells, most of the EGFR signal was detected together with the actin-rich filopodia at the cell cortex, whereas only a small amount of EGFR staining was detected inside the cells (**Fig. 6A**). In A431 KO cells, on the other hand, most of the EGFR staining was observed in intracellular punctae (**Fig. 6B**). Orthogonal xz-projections through the cells revealed that most of the EGFR staining in A431 control cells overlapped with the cortical actin signal, while in A431 KO cells part of the signal was detected inside the cells and a fraction near the cortex underneath the cortical actin (**Fig. 6A’-B’’**). Moreover, A431 KO cells appeared significantly flatter than A431 control cells.

**Figure 6.**
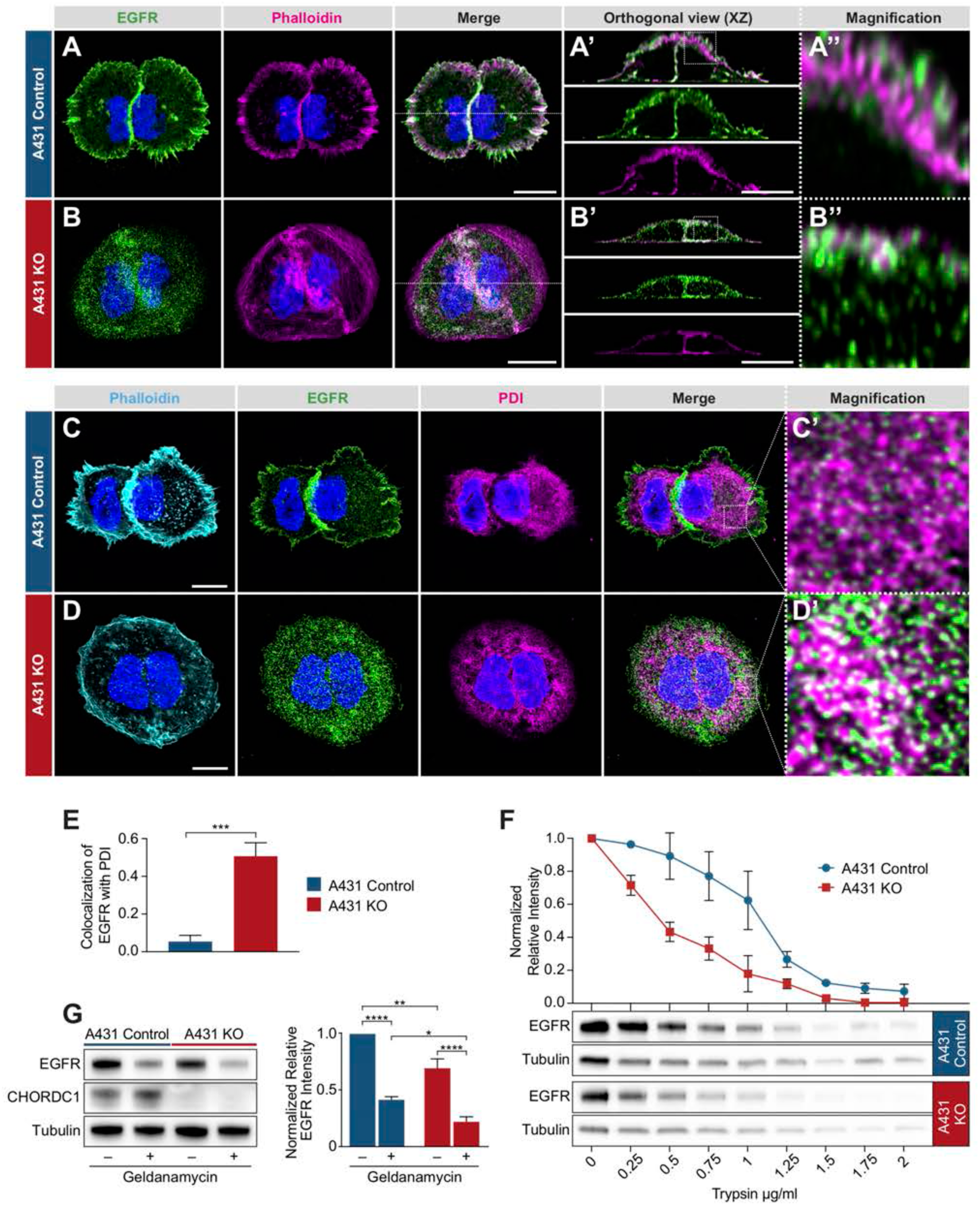
Mislocalization of the EGFR in CHORDC1 mutant A431 cells. (**A**) Immunofluorescence staining of A431 control and (**B**) A431 KO cells with antibodies recognizing EGFR (green), fluorescently-labeled phalloidin (magenta) and DAPI (blue). The dotted lines in the merge panels indicate the planes used to create the orthogonal (XZ) views shown in (**A’**, **B’**). The doted squares in (**A’**, **B’**) indicate the areas shown at higher magnification in (**A’’**, **B’’**). (**C**) Immunofluorescence staining of A431 control- and (**D**) KO cells with fluorescently-labeled phalloidin (light blue), antibodies recognizing EGFR (green), the endoplasmic reticulum marker PDI (magenta), and with DAPI (blue). (**C’**, **D’**) show higher magnifications of the regions outlined with dotted squares in the merge panels of (**C, D**). All images are maximum intensity projection of three confocal sections. The scale bars are 10 µm. (**E**) Co-localization of EGFR and PDI in A431 control (n=5) and A431 KO cells (n=10) was quantified by calculating the Mander’s coefficient as described in materials and methods (Manders *et al*, 2011). p-values were calculated by a two-tailed t-test for independent samples. Error bars show the SEM. (**F**) Trypsin sensitivity assay. Total protein extracts of A431 control and A431 KO cells were incubated with the indicated trypsin concentrations, and the samples were analyzed by Western blotting with antibodies against EGFR and tubulin. The line graph shows a quantification of the EGFR levels double normalized to the tubulin signal in each sample and to the undigested (0 µg/ml) samples. Error bars show the SEM. The average of three biological replicates is shown. (**G**) Western blot analysis of EGFR and CHORDC1 protein levels in A431 control and A431 KO cells with and without 1 µM geldanamycin treatment. The bar graph shows the normalized averaged relative intensities ± SEM. p-values (p < 0.05 = *, p < 0.01 = ** and p < 0.0001 = ***) were calculated by one-way ANOVA and corrected with a Tukey multiple comparison test. The average of three biological replicates is shown.

To examine if the loss of CHORDC1 results in a similar mislocalization of the EGFR to the ER as in *C. elegans chp-1(lf)* mutants, we co-stained the cells with antibodies against the EGFR and the ER marker PDI (protein disulfide isomerase) (Jaronen *et al*, 2013). While in A431 control cells only 5% of the EGFR signal colocalized with the PDI marker, 51% of the EGFR signal in the A431 KO cells overlapped with the PDI staining (**Fig. 6C-E**).

Since CHORDC1 has been reported to act as a co-chaperone (Gano & Simon, 2010a), we hypothesized that loss of CHORDC1 function might result in the incorrect folding of the EGFR and thereby cause its accumulation in the ER. To test this hypothesis, we performed trypsin sensitivity assays of the EGFR in A431 control versus KO cells. Since misfolded proteins are usually more susceptible to trypsin degradation, this assay can detect changes in protein folding (Ninagawa *et al*, 2015). Total protein extracts of A431 control and KO cells were incubated with varying concentrations of trypsin, and the amount of full-length EGFR was quantified by Western blotting. As internal control, the EGFR signal intensities were normalized to the tubulin levels in the same lysates, as tubulin was degraded at approximately the same rate in the two cell populations. This assay revealed a faster degradation and thus an increased trypsin sensitivity of the EGFR in A431 KO cells(**Fig. 6F**).

Consistent with earlier reports (Ahsan *et al*, 2012), we observed that the inhibition of HSP90 by geldanamycin caused an approximately two-fold reduction in total EGFR levels (**Fig. 6G**). By contrast, loss of CHORDC1 (A431 KO) cells only resulted in a 20% reduction in EGFR levels. Surprisingly, geldanamycin treatment of A431 KO cells caused a further reduction below the EGFR levels detected in geldanamycin-treated A431 control cells, indicating that HSP90 stabilizes the EGFR independently of CHORDC1.

Taken together, we have found that the EGFR accumulates in the ER of A431 cells lacking CHORDC1, analogous to the mislocalization of LET-23 in *C. elegans chp-1(lf)* mutants. The increased trypsin sensitivity of the mislocalized EGFR in CHORDC1 mutant cells could be due to incorrect protein folding. Notably, CHORD1 and HSP90 appear to influence EGFR stability through distinct mechanisms.

## Discussion

Intercellular signal transduction is regulated by the production and secretion of growth factors in the signal sending cells, as well as by the sub-cellular localization and intracellular trafficking of their receptors in the signal receiving cells (Sorkin & Goh, 2009). We have used vulval development in *C. elegans* as an in vivo model to identify new factors regulating the secretion and localization of the EGFR homolog LET-23 (Haag *et al*, 2014). The basolateral localization and retention of LET-23 is essential for the efficient activation of the downstream RAS/MAPK pathway and correct VPC differentiation. Perturbations in LET-23 secretion or localization invariably cause defects in VPC fate specification and abnormal vulval morphogenesis.

In a screen for genes regulating LET-23 localization we have previously identified the CHORD-containing protein CHP-1 as a regulator of LET-23 EGFR trafficking in the VPCs (Haag *et al*, 2014). Here, we show that the loss of *chp-1* function in *C. elegans* leads to the accumulation of LET-23 in the ER and a strong reduction -but not a complete inactivation- of RAS/MAPK signaling in the VPCs. Even in the absence of CHP-1 a small fraction of LET-23 reaches the plasma membrane, where it can bind to and be activated by the normally limiting amounts of LIN-3 EGF. Our genetic analysis confirms the notion that vulval induction in *chp-1(lf)* mutants depends to a large extent on *lin-3* activity. However, the ER accumulation of LET-23 in *chp-1(lf)* mutants is independent of *lin-3* or the basolateral *lin-2/lin-7/lin-10* receptor localization complex. This indicates that CHP-1 controls LET-23 secretion in the VPCs at an earlier step, before the receptor interacts with LIN-7 at the plasma membrane and undergoes LIN-3-mediated endocytosis.

In order to test if the function of CHP-1 in the EGFR signaling pathway is conserved in mammals, we inactivated the CHP-1 homolog CHORDC1 in human A431 epidermoid carcinoma cells, which express high levels of the wild-type EGFR and undergo a phenotypic switch in response to EGF stimulation (Ferretti *et al*, 2011; Van de Vijver *et al*, 1991). The phenotype of the CHORDC1 knock-out in A431 cells is remarkably similar to the *chp-1(lf)* phenotype in the *C. elegans* VPCs; the EGFR accumulates in the ER and the activation of the RAS/MAPK pathway in response to EGF stimulation is strongly reduced. Even though the total levels of the EGFR are only slightly reduced CHORDC1 mutant A431 cells, the EGFR is less stable as it exhibits an increased sensitivity to trypsin digestion (Ninagawa *et al*, 2015). Possibly, CHORDC1 is required for the correct folding of the EGFR as it enters the ER and the partially unfolded EGFR molecules cannot pass through the ER. Our current data do not distinguish whether CHORDC1 is required for the ER entry or exit of the EGFR. However, the extracellular domain of the EGFR is N-glycosylated at multiple sites after entry into the ER lumen and further modified in the Golgi network to produce a mature glycoprotein with a molecular weight of 170 Kilodaltons (kDa), as opposed to the 130 kDa native, non-glycosylated polypeptide (Azimzadeh Irani *et al*, 2017). Since we did not observe a shift in the electrophoretic mobility of the EGFR in A431 KO cells, it appears that the EGFR is at least partially glycosylated and hence can enter the ER without CHORDC1.

Several observations have indicated that the role of *C. elegans* CHP-1 in LET-23 EGFR trafficking is rather specific. First, if CHP-1 was required for the ER trafficking of a large number of proteins, this would result in ER stress and activate the UPR pathway (Calfon *et al*, 2002). Yet, *chp-1(lf)* mutants do not exhibit an increased activity of the UPR pathway, unless additional ER stress is induced by globally inhibiting protein glycosylation. Second, the membrane localization of three other type I trans-membrane receptors we examined (LIN-12 NOTCH, PAT-3 ß-integrin and LIN-18 RYK) does not depend on *chp-1*. Third, inhibition of other known HSP90 co-chaperones, such as *cdc-37*, *daf-41* and *sgt-1*, does not affect LET-23 localization in the VPCs.

It has been proposed that CHORDC1 acts as a co-chaperone that assists HSP90 in the folding of its numerous client proteins (Gano & Simon, 2010a). According to this model, CHORDC1 would confer the specificity of HSP90 towards a subset of its clients, among them the EGFR. Surprisingly, our data point at an HSP90-independent function of CHP-1/ CHORDC1 in EGFR trafficking. The strongest viable allele of *daf-21*, which encodes the *C. elegans* HSP90 ortholog, did not perturb LET-23 membrane localization, though the expression levels of LET-23 in the VPCs of *daf-21(rf)* mutants were reduced. On the other hand, the vulval cells of *chp-1(lf)* mutants did not exhibit an obvious reduction in LET-23 expression. We made analogous observations using human A431 cells. The treatment of A431 cells with the HSP90 inhibitor geldanamycin caused a strong reduction in total EGFR protein levels, while loss of CHORDC1 only caused a slight (20%) reduction in total EGFR levels. Since geldanamycin treatment reduced EGFR levels even in CHORDC1 deficient cells, CHORDC1 likely functions independently of HSP90. The CDC37/HSP90 co-chaperone/chaperone complex interacts with the nascent EGFR and may assists in its folding (Verba & Agard, 2017), whereas CHORDC1 might promote the maturation and trafficking of EGFR independently of HSP90. Furthermore, many co-chaperones can act independently of their core chaperones. For example, p23, which contains the same ACD (alpha-crystallin-Hsps_p23-like) domain as CHORDC1, regulates various cellular processes that are distinct from those controlled by HSP90 (Echtenkamp et al, 2011). It is therefore possible that CHORDC1 acts by itself or in a complex with another member of the large heat shock protein family. For example, the glucose-regulated protein 94 (GRP94), an HSP90 paralog localized in the ER, regulates the intracellular trafficking of Toll-like receptors (Randow & Seed, 2001). It thus remains to be tested whether CHORDC1 acts in a complex with GRP94 or other chaperones.

A number of clinical trials have tested HSP90 inhibitors for the treatment of human cancer, geldanamycin being among the first generation of HSP90 inhibitors used (Garcia-Carbonero *et al*, 2013). However, due to the large number of HSP90 clients the use of these inhibitors in patients has been limited by the many side effects they cause. The identification of CHORDC1 as a more specific regulator of EGFR trafficking opens the possibility to develop small molecule chaperone inhibitors targeting the EGFR pathway with a higher selectivity and fewer side effects.

## Materials and Methods

### General *C. elegans* methods and strains

Unless specified otherwise, *C. elegans* strains were maintained at 20 °C on Nematode Growth Medium (NGM) agar plates as described (Brenner, 1974). The *C. elegans* Bristol N2 strain was used as wild-type reference, and all strains generated through genetic crosses were derived from N2. A complete list of the *C. elegans* strains used can be found in **suppl. Table S1**.

### RNAi feeding method

RNAi feeding experiments were performed as described previously (Kamath & Ahringer, 2003). The strain of interest was fed with *E. coli* HT115 expressing dsRNA against a specific target mRNA. 20 synchronized L1 larvae were transferred to NGM plates containing 3 mM IPTG and 50ng/ml ampicillin seeded with the indicated RNAi bacteria. The F1 progeny of the 20 P0 animals was analyzed at the L3 stage to score LET-23::GFP localization or at the L4 stage to examine vulval induction.

### Vulval induction

Vulval induction was scored by examining 20-40 worms of the indicated genotypes at the L4 stage under Nomarski optics. Animals were mounted on 4% agarose pads and anesthetized with 20 mM tetramisole in M9 buffer as described (Sternberg & Horvitz, 1986; Schmid *et al*, 2015). The vulval induction index (VI) was scored by counting the induced VPCs in 20-40 animals and calculating the average number of induced VPCs per animal. Statistical analysis is described in the legend to **Fig. 4**.

### Tunicamycin treatment

The tunicamycin treatment to induce ER stress was carried out as described in (Taylor & Dillin, 2013). Briefly, animals expressing the *hsp-4::gfp* reporter were synchronized at the L1 stage and allowed to develop on NGM plates until the first day of adulthood. Then, they were incubated for 4 hours at room temperature in 25 µg/ml tunicamycin solution in M9 buffer. Control animals were incubated in an equivalent dilution of DMSO, which was used as a solvent for tunicamycin. After the treatment, HSP-4::GFP expression was observed with a 10x lens on a Leica DM RA wide-field microscope. Images were analyzed using Fiji software (Schindelin *et al*, 2012), and the average intensity of the whole body in each animal was measured to make the box plot in **suppl. Fig S3**.

### Generation of the SP12 ER and AMAN-2 Golgi reporters

Plasmid constructs were made using Gibson Assembly cloning (Gibson *et al*, 2009) and verified by DNA sequencing. A list of the oligonucleotide primers used for plasmid construction can be found in **suppl. Table S2**. To construct plasmid pAHE3 (P*dlg-1::aman-2::mCherry::unc-54 3‘UTR,* the plasmid the pCFJ151 backbone (Frøkjær-Jensen *et al*, 2008) was recombined with the P*dlg-1* promoter and the *unc-54* 3’ UTR, amplified as two individual fragments using the primers OEH153& OEH158 and OEH159& OEH156. The *aman-2* (F58H1.1) genomic sequence encoding the first 82 amino acids, including the signal sequence and trans-membrane anchor, amplified from genomic DNA using the primers OEH152& OEH155, and the mCherry coding sequence, amplified with the primers OEH154& OEH157, were then inserted after the P*dlg-1* promoter.

To construct pAHE6 (P*dlg-1::mCherry::C34B2.10(SP12)::unc-54 3‘UTR),* the pCFJ151 backbone (Frøkjær-Jensen *et al*, 2008) was recombined with the P*dlg-1* promoter and the *unc-54* 3’ UTR, amplified as two individual fragments using the primers OAHE22& OEH159 and OEH158& OEH153. The mCherry coding sequence, amplified with the primers OAHE19 & OAHE20, and the genomic sequence of C34B2.10 (SP12) containing the stop codon, amplified with the primers OAHE21& OAHE8, were then inserted after the P*dlg-1* promoter. The primer OAHE21 contained an additional linker sequence encoding three Alanines for the N-terminal fusion with mCherry. For each of the two reporter plasmids, single copy insertion transgenes were generated by the MosSCI method as described (Frøkjær-Jensen *et al*, 2008).

### VPC-specific *chp-1* CRISPR/CAS9

Two target sites in the first exon of the genomic *chp-1* locus (sgRNA #1: CAG TGC TAT CAT AAA GGA TG and sgRNA #2 CGG TCT CCT TTT CGA TCC CA) were identified using the CRISPR design tool (http://crispr.mit.edu/) (Hsu *et al*, 2013). Double-stranded oligonucleotides were synthetized and cloned into the pDD162 (Addgene) to produce pAHE4 (sgRNA #1) and pAH5 (sgRNA #2). Plasmid pEV5 (*Pegl-17-Δpes-10::cas9*) (gift by Evelyn Lattmann) was used for 1° VPC-specific expression of the CAS9 protein. The plasmids pAHE4 and pAHE5 were co-injected into the gonads of wild-type animals as described (Mello *et al*, 1991) at a concentration of 50 ng/μl each together with the plasmid pEV5 at 100 ng/μl and the transformation markers pGH8 (P*rab-3::mCherry*) at 10 ng/ μl, pCFJ104 (P*myo-3::mCherry*) at 5 ng/μl and the pCFJ90 (P*myo-2::mCherry*) at 2.5 ng/μl to create the extrachromosomal array *zhEx558*.

### Mammalian Cell Culture

The human vulva epidermoid carcinoma cell line A431 was obtained from Sigma Aldrich (85090402) and cultured in Dulbecco’s Modified Eagle Medium (Gibco 41966-029) according to standard mammalian tissue culture protocols and sterile technique. DMEM was supplemented with 10% FCS (Gibco 10500-064) and 1% Pen-Strep (Gibco 15140-122).

### CHORDC1 knock-out in A431 cells

CHORDC1 guide RNAs targeting the first exon of CHORDC1 (CHORDC1 sgRNA #1: TTA CCG TCG GAA TTG GTC TC and CHORDC1 sgRNA #2: AGA CCA ATT CCG ACG GTA AG), as well as the scramble sequence GCA CTA CCA GAG CTA ACT CA, were identified using the CRISPR Design Tool (http://crispr.mit.edu/) (Hsu *et al*, 2013). Double-stranded oligos were generated and cloned into the lentiCRISPRv2 vector (Addgene), which was then transfected in combination with pVSV-G, pMDL and pREV into HEK293T cells to produce lentiviral particles (vMW6_CHORDC1 sg#1, vMW7_CHORDC1 sg#2, vMW9_scramble). Four days following transfection, the media from cells was collected, clarified by centrifugation, and filtered through a 0.45 μM filter to collect lentiviral particles. Subsequently, the particles were concentrated in Amicon Ultra tubes (Ultracel 100k, Millipore). The titer of viral particles was determined before vMW6_CHORDC1 sg#1 and vMW7_CHORDC1 sg#2 were used to transduce 180’000 A431 cells in a 12-well plate at a combined MOI of 10. A431 cells were supplemented with DMEM media containing the lentiviral mix and 10 μg/ml polybrene and cultured under normal conditions. In parallel, cells were transduced with vMW9_scramble at a MOI of 10. Three days after transduction, cells were grown in the presence of 1.2 μg/ml puromycin. One week after puromycin selection, the puromycin-resistant populations were frozen and kept as stocks used in the subsequent experiments.

### Western blotting

For Western blot analysis, cells were lysed in lysis buffer on ice (100 mM Tris/HCl, 150 mM NaCl, 1 % Triton X, 1 mM EDTA, 1mM DTT, 10 ml lysis buffer + 1 tablet protease inhibitor), scraped with a cell scraper and snap frozen in liquid nitrogen. 100 µl of this mix was sonicated in a Bioruptor sonicator device (Diagenode), before 100 µl of 2x SDS loading dye were added. About 10 μg of protein extract were resolved by SDS-PAGE and transferred to nitrocellulose membrane. The membrane was blocked in 5% dried milk in 1x PBS plus 0.2% Tween 20 and then incubated with the diluted primary antibodies overnight at 4°C. Secondary anti-rabbit or anti-mouse IgG antibodies conjugated to horseradish peroxidase (HRP) were used as the secondary antibodies. The HRP was detected by incubating the membrane with the SuperSignal West Pico or Dura Chemiluminescent Substrate (Thermo Scientific) for 4 minutes, before the signals were measured on a digital Western blot imaging system. The antibodies used for Western blot analysis were: anti-CHORDC1 (HPA041040 Atlas Antibodies), anti-Tubulin (ab18251 abcam), anti-EGFR (HPA018530 Atlas Antibodies), anti-ERK1/2 (M5670 Sigma Aldrich), anti-ERK1/2 activated (M8159, Sigma Aldrich), HRP anti-Rabbit (111-035-144 Jackson Immuno Research) and HRP anti-Rabbit (115-035-146 Jackson Immuno Research). Quantification of Western blots was done by measuring the band intensities in Fiji.

### Immunofluorescence staining of A431 cells

Cells were grown on glass slides in 24-well plates under standard conditions for 48 hrs. Slides were then rinsed in PBS and fixed for 15 min in 4% PFA at 37°C. After washing with PBS, cells were permeabilized with PBS containing 0.2% Triton X-100 and 0.5% BSA for 5 min, and then blocked for 1 hour in PBS containing 0.5% BSA and 0.2% gelatine. Primary antibodies were added in blocking solution for 1 hour at room temperature in a humid chamber. The cells were rinsed in PBS three times before being incubated for 40 minutes in the dark with secondary antibodies and Phalloidin 568 (B3475 Thermo Scientific). After three washes with PBS, cells were stained for 5 minutes with PBS containing 0.1 µg/ml DAPI, followed by three washes with PBS. Glass slides were mounted with ProLong Gold Antifade Mountant (Thermo Scientific). The antibodies used for immunocytochemistry were: anti-CHORDC1 (HPA041040 Atlas Antibodies), anti-Tubulin (ab18251 abcam), anti-EGFR (MA5-13269 Thermo Scientific), anti-PDI (MA3-019 Thermo Scientific), Alexa Fluor 488 (A11034 Thermo Scientific), and Alexa Fluor 647 (A21236 Thermo Scientific). For all antibody stainings, at least three biological replicates were made.

### EGF stimulation

250’000 A431 cells were grown in 12-well plates under standard conditions for 24 hrs. Thereafter, growth medium was replaced with DMEM lacking FCS. After 15 hours of starvation, cells were stimulated with 100 ng/ml human EGF (E9644 Sigma) for 10 minutes at 37°C, before they were lysed and prepared for western blot analysis.

### Trypsin Sensitivity Assay

750’000 A431 cells were seeded into a 25cm^2^ flask and grown until they reached confluency. After washing twice with cold PBS, cells were scraped with a cell scraper, lysed in lysis buffer (100mM Tris pH=8, 1% NP-40, 150 mM NaCl, 1 mM DTT) for 10 min on ice, sonicated for 10 min at 4°C, and clarified by centrifugation at 17,000 x g for 10min at 4°C. Aliquots containing 50 µg of cleared protein samples were incubated with 0.1, 0.15, 0.2, 0.4, 0.6, 0.8, 1, 1.25 and 1.5 µg/ml Trypsin (Sigma EMS0004) and incubated for 15 min at 25°C while shaking. Thereafter, 2x SDS loading dye was added and samples were boiled prior to resolving the proteins by SDS-PAGE.

### Wide-field fluorescence microscopy

To examine the expression pattern of fluorescently tagged proteins, a Leica DM RA wide-field microscope equipped with a Hamamatsu ORCA-ER camera using a 40x /1.3 NA or 63x/1.4 NA oil immersion objective was used. Fluorescent and Nomarski mages were acquired with the Openlab 4.0 or VisiView 2.1 software. To observe the localization of GFP reporters, around 40 worms at the L3 stage were mounted on 4% agarose pads and anesthetized with 20 mM tetramisole in M9 buffer. In each experiment, the intensity of the light source, the exposure time and the software settings were kept constant.

### Confocal laser scanning microscopy

Around 40 larvae at the Pn.p to Pn.pxx stage were mounted on 4% agarose pads in M9 buffer containing 2 mM tetramisole. z-stacks at 0.2 to 0.3 μm spacing were taken using a Plan-Apochromat 63x/1.4 NA oil immersion objective on a Zeiss LSM710 confocal laser scanning microscope equipped with an 458/488/514 nm argon laser and a 594 nm helium-neon laser. Images were acquired by using the LSM710 ZEN 2012 software (Zeiss). GFP was excited with a 488 laser excitation and emission was detected in a range of 493-566 nm. mCherry was excited with a 594 nm laser excitation and emission was detected in a range of 599-696 nm. Images were captured with a variable frame size, a pinhole equivalent to 1 Airy and a pixel size of 0.08 μm. Identical camera gain settings were used for all live animal imaging (AMAN-2::mCherry: GFP 600, mCherry 500/ mCherry::SP12: GFP 600, mCherry 600). Images of antibody-stained A431 cells (**Figs. 5 & 6**) were taken on a Leica CLSM SP8 upright microscope equipped with 405/488/552/638 nm solid state diode lasers. z-stacks at 0.16 µm spacing were taken using a 63x/1.4 HCX PL APO CS2 oil immersion objective with a variable frame size and a pinhole equivalent to 1 Airy.

### Image processing Quantification of co-localization

Images were analyzed and processed with Fiji software (Schindelin *et al*, 2012) to adjust brightness and contrast. Images in **Figs. 2, 5 & 6** were deconvolved using the Huygens deconvolution software (Scientific Volume imaging). To quantify co-localization in **Figs. 2 & 6**, raw images (without deconvolution) were processed in Fiji using the Subtract Background command (default settings, rolling ball radius=0.3). To quantify co-localization, z-stacks were analyzed with the ImarisColoc module in Imaris 8.3. (Bitplane). The GFP channel of each image was thresholded to define a ROI using the masking channel function. An automatic threshold implemented in Imaris was set for both channels based on a statistical significance algorithm (Costes *et al*, 2004). With this approach, the extent of co-localization of two fluorescent-labeled proteins in an image is automatically quantified, without the bias of visual interpretation. Based on an automatically identified threshold calculated by the Coste’s approach, a Manders Colocalization Coefficient (MCC) indicating the fraction of total probe fluorescence that co-localizes with the fluorescence of a second probe was calculated (Manders *et al*, 2011). The calculated MCC values were averaged, and statistical analysis was performed using a a two-tailed Student’s t-test for independent samples.

## Acknowledgements

We wish to thank the members of the Hajnal laboratory for critical discussion and comments on the manuscript. We are also grateful to the *C. elegans* Genetics Center CGC, which is funded by NIH Office of Research Infrastructure Programs (P40 OD010440), the Mitani lab (National Bioresource Project) for providing some strains, Andrew Fire for GFP vectors, J. Ahringer for RNAi clones and Franziska Walser and Marco Wachtel for their expertise in the production of lentiviral particles. Confocal imaging was performed with support of Urs Ziegler at the Center for Microscopy and Image Analysis, University of Zurich. This work was supported by a grant from the Swiss National Science Foundation to A.H. no. 31003A-166580 and the Kanton of Zürich.

The authors declare no competing interests.

## Supplementary information

**suppl. Fig. S1.**
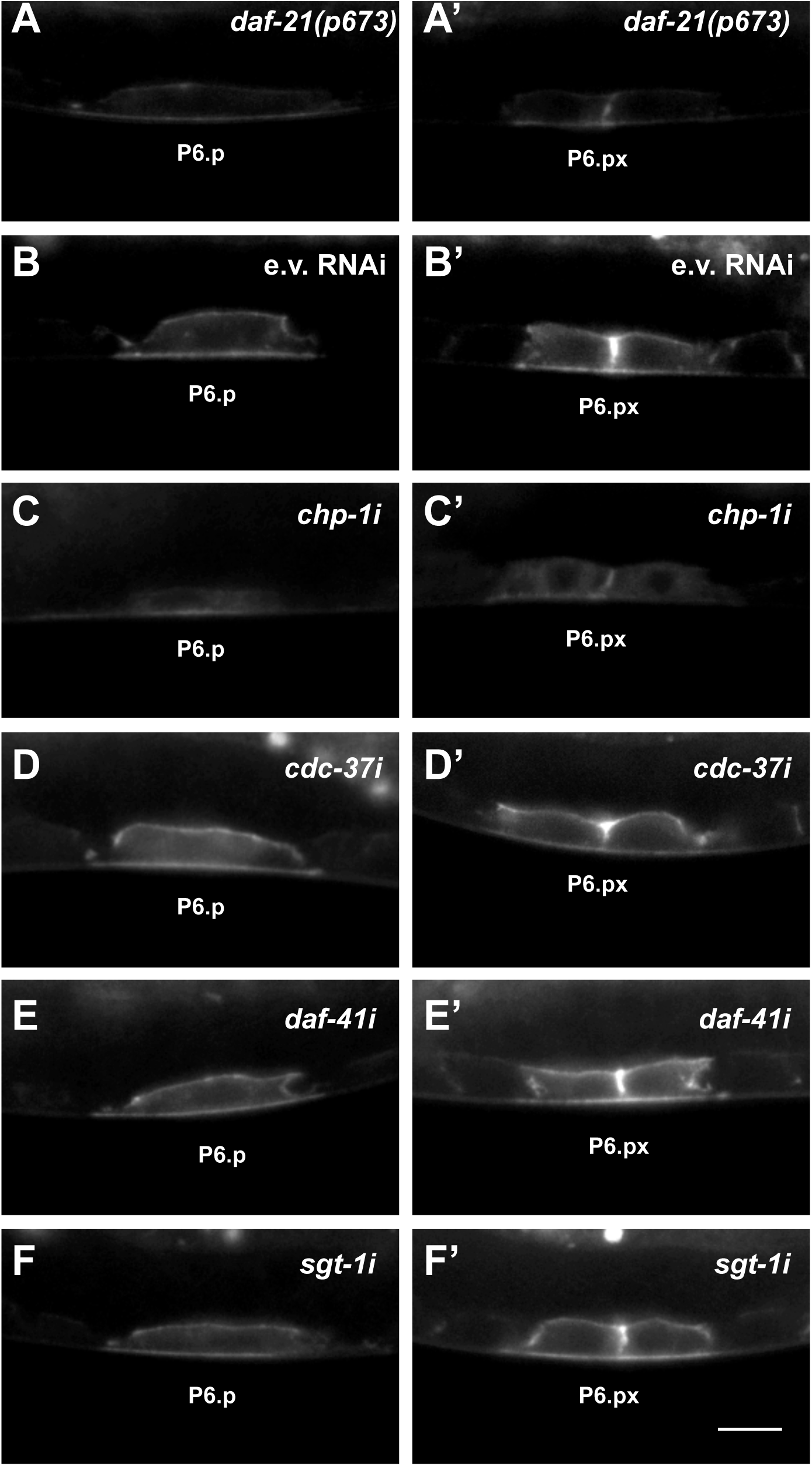
LET-23::GFP localization in *daf-21(p673)* mutants and after RNAi knock-down of co-chaperones. LET-23::GFP localization in P6.p and (left panels) the two P6.p daughters (right panels) in (**A,A’**) *daf-21(p673)* mutants and (**B-F’**) under the indicated RNAi conditions. E.v. RNAi in (**B,B’**) are the negative controls treated with empty RNAi vector. At least 20 animals were analyzed for each condition.

**suppl. Fig. S2.**
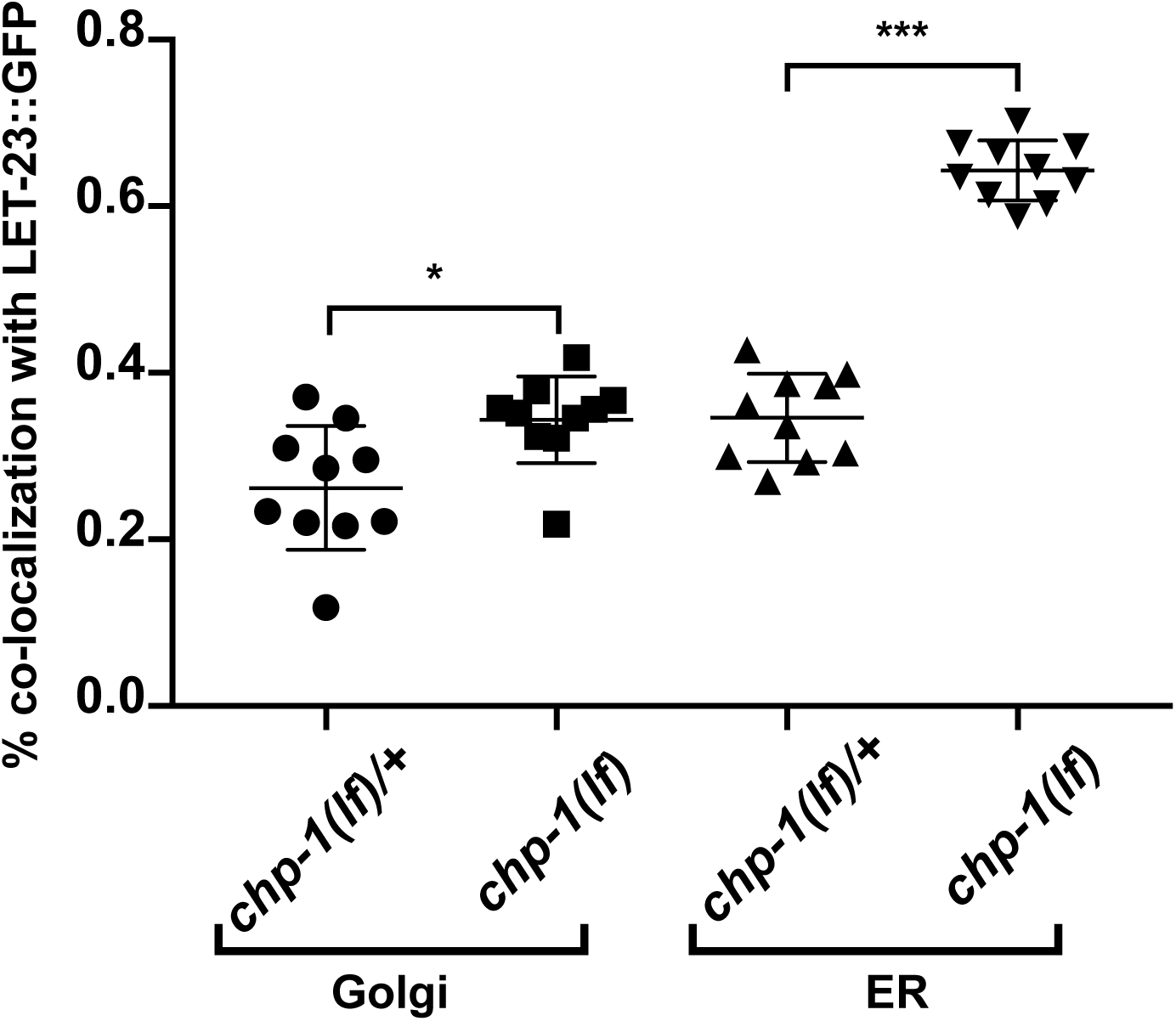
Quantification of the co-localization between the AMAN-2::mCherry (Golgi) and SP12::mCherry (ER) markers with LET-23::GFP in P6.p and its descendants. Co-localization of the AMAN-2::mCherry Golgi or the SP12::mCherry ER marker with LET-23::GFP was quantified by calculating the thresholded Mander’s coefficient (MANDERS *et al*, 2011) as described in materials and methods. Error bars show the standard error of the mean (SEM). p-values were calculated by two-tailed t-tests for independent samples (p < 0.05 = * and p < 0.001 = ***). Ten animals were analyzed for each condition.

**suppl. Fig. S3.**
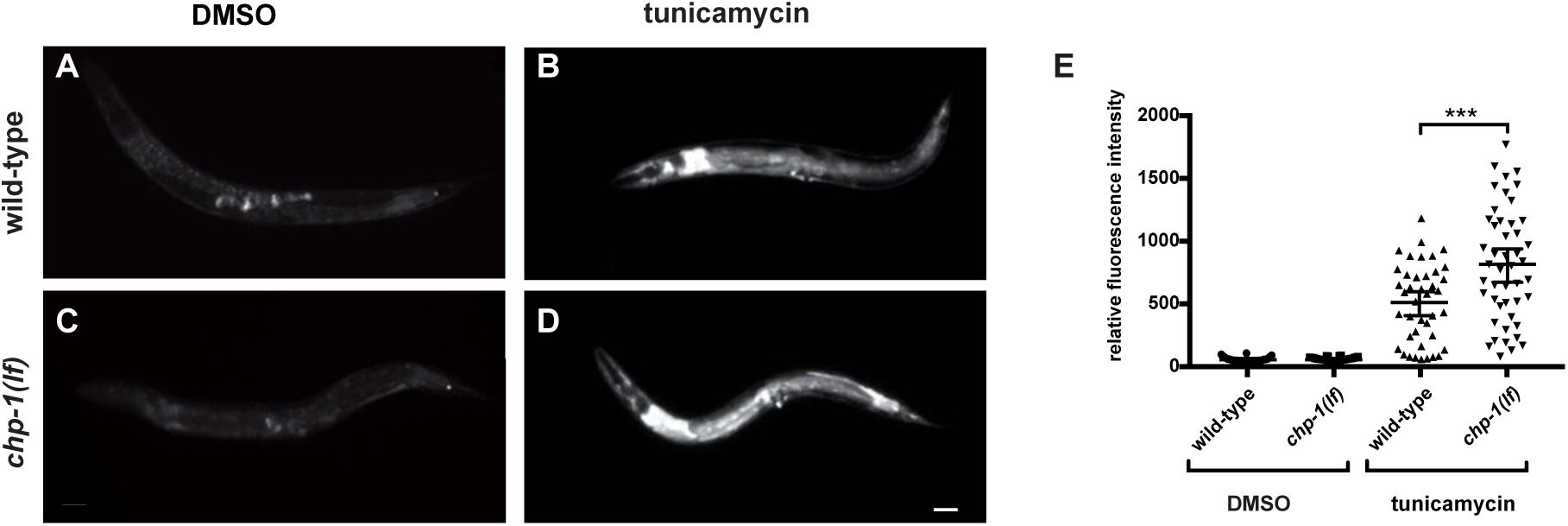
Unfolded protein response in *chp-1(tm2277lf)* mutants after tunicamycin treatment. (**A**) HSP-4::GFP expression in untreated (DMSO) controls and (**B**) tunicamycin treated wild-type animals, and (**C**) in untreated and (**D**) tunicamycin treated *chp-1(lf)* mutants. The scale bar is 10µm. (**E**) Quantification of HSP-4::GFP signal intensities under the different conditions. Error bars indicate the 95% CI. p-values were calculated by t-tests for independent samples and are indicated as *** for p<0.01. Between 26 and 47 animals were analyzed for the different conditions.

**suppl. Table S1.**
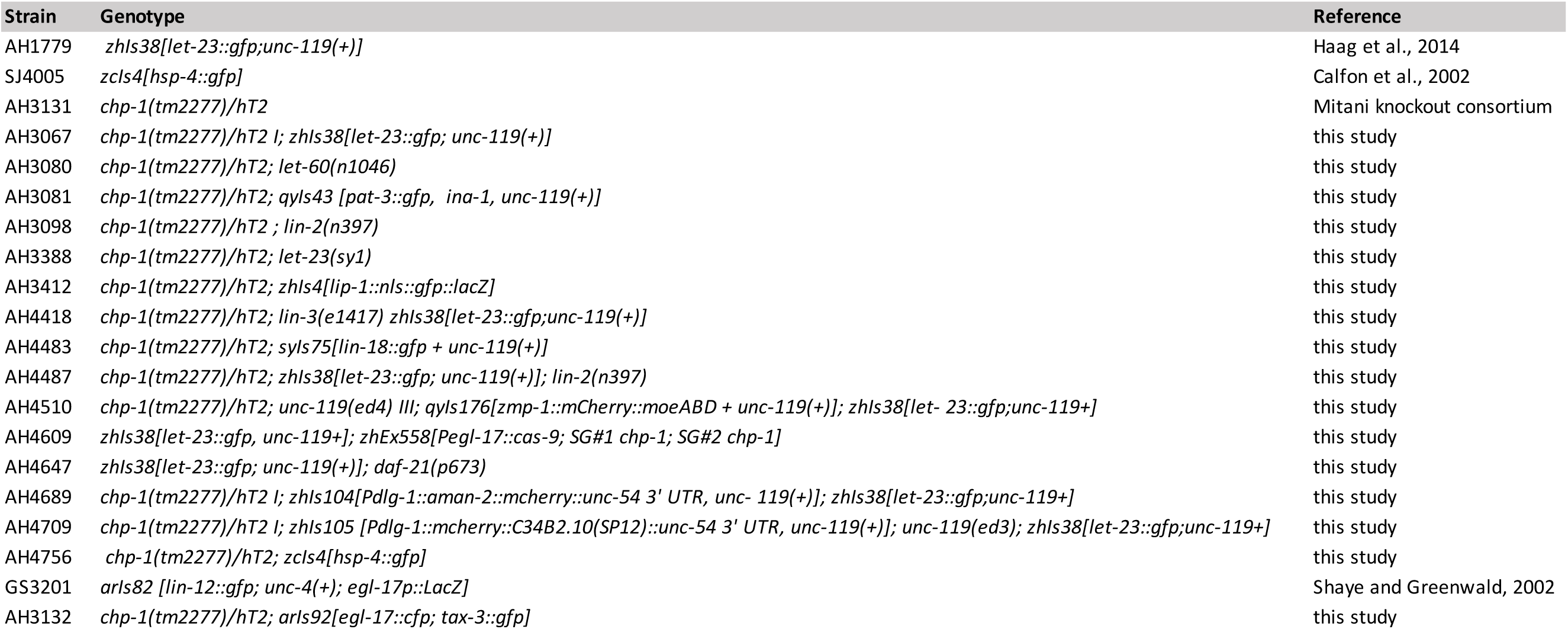
Genotypes of the strains used in this study.

**suppl. Table S2.**
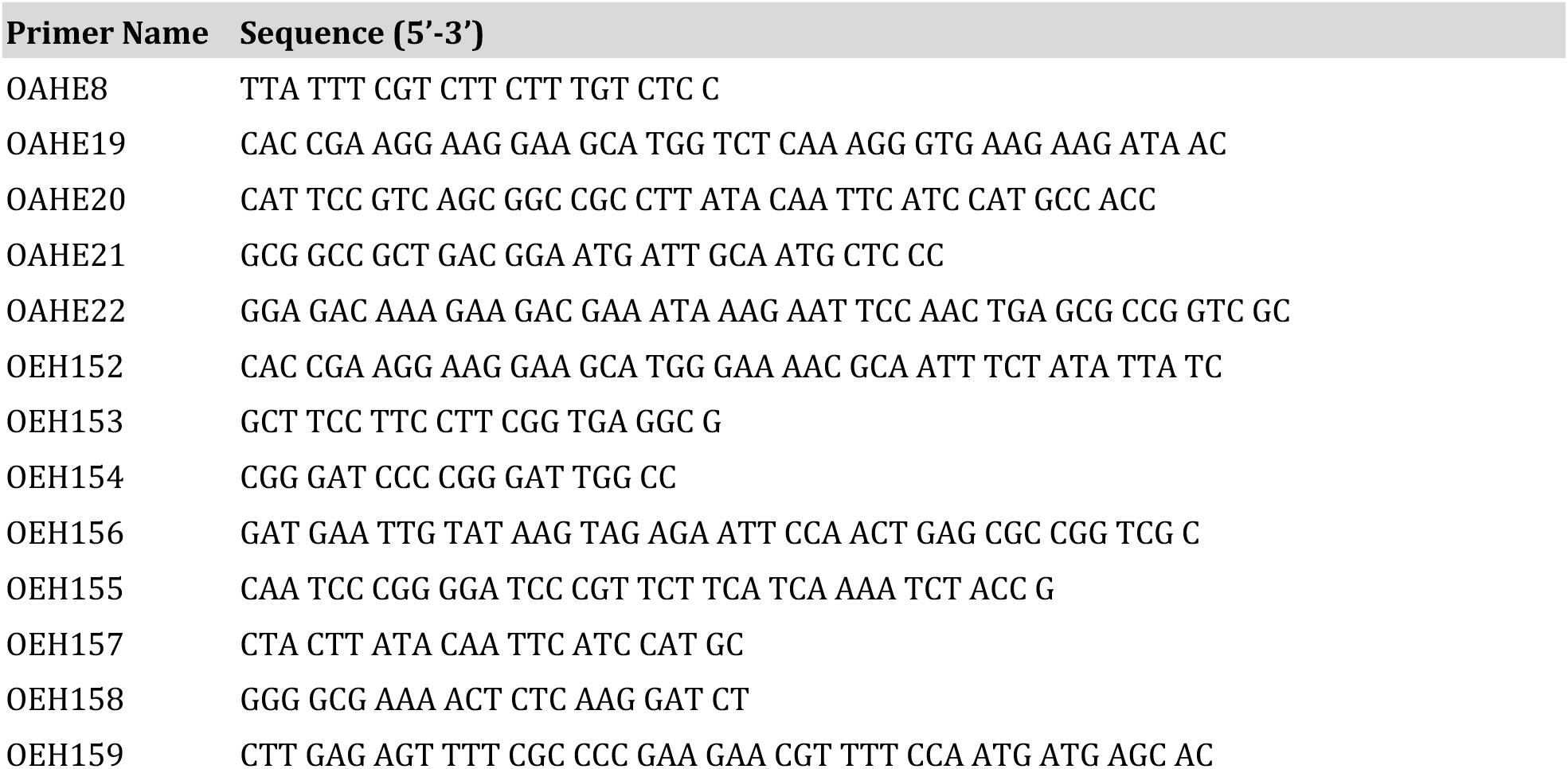
Oligonucleotide primers used for plasmid constructions.

